# Quantifying cell densities and biovolumes of phytoplankton communities and functional groups using scanning flow cytometry, machine learning and unsupervised clustering

**DOI:** 10.1101/274357

**Authors:** Mridul K. Thomas, Simone Fontana, Marta Reyes, Francesco Pomati

## Abstract

Scanning flow cytometry (SFCM) is characterized by the measurement of time-resolved pulses of fluorescence and scattering, enabling the high-throughput quantification of phytoplankton morphology and pigmentation. Quantifying variation at the single cell and colony level improves our ability to understand dynamics in natural communities. Automated high-frequency monitoring of these communities is presently limited by the absence of repeatable, rapid protocols to analyse SFCM datasets, where images of individual particles are not available. Here we demonstrate a repeatable, semi-automated method to (1) rapidly clean SFCM data from a phytoplankton community by removing signals that do not belong to live phytoplankton cells, (2) classify individual cells into trait clusters that correspond to functional groups, and (3) quantify the biovolumes of individual cells, the total biovolume of the whole community and the total biovolumes of the major functional groups. Our method involves the development of training datasets using lab cultures, the use of an unsupervised clustering algorithm to identify trait clusters, and machine learning tools (random forests) to (1) evaluate variable importance, (2) classify data points, and (3) estimate biovolumes of individual cells. We provide example datasets and R code for our analytical approach that can be adapted for analysis of datasets from other flow cytometers or scanning flow cytometers.

## Introduction

Flow cytometry (FCM) has enabled the monitoring of natural microbial communities by capturing point estimates of cellular characteristics and images, respectively ([1–7]. Developed more recently, scanning flow cytometry (SFCM) records time-resolved pulses of fluorescence and scattering for every cell [4,8–10]. The fluorescence and scattering pulses are summarized using parameters that characterise changes in morphology and pigmentation over the length of a cell. This vast amount of individual-level information can be used to quantify the distributions of important cellular *traits* within communities. High-throughput quantification of traits governing organism-environment interactions [11] would increase our ability to understand ecological and evolutionary changes in microbial communities. However, the utility of traditional FCM and scanning flow cytometry (SFCM) in monitoring natural communities has been limited by a lack of protocols that allow us to automate 1) cleaning of SFCM data by removing signals that do not belong to living cells, 2) classifying individual cells into functional groups, and 3) quantifying the biovolumes of individual cells, and the total biovolumes of the whole community and major functional groups (we follow convention by referring to cyanobacteria, diatoms, etc. as ‘functional groups’, but these groups are defined taxonomically). The first two goals are a challenge in FCM and SFCM analyses because validating results from these analyses is difficult in the absence of images (which are available only in imaging flow cytometers, for which high-throughput analytical methods are available e.g. [6,12]). This paper demonstrates a protocol to achieve these three goals in FCM or SFCM datasets.

Cleaning datasets (i.e. removing signals that are not from live cells) and identifying functional groups in FCM has traditionally been done manually. This has involved visually identifying clusters of points with similar trait values using small numbers of bivariate plots. These manually-identified clusters are used to separate measurements of live cells from other signals (‘gating’), and to identify distinct cell types for further analyses. However, a manual sample-by-sample approach is not practical when large numbers of samples are measured, such as in high-frequency monitoring efforts or large experiments. Additionally, the large number of variables of unknown importance captured by SFCM makes the choice of variables for plotting difficult, and the results sensitive to these choices. Previous efforts to avoid these problems in FCM and SFCM have involved focusing on very small numbers of variables and using (i) linear thresholds for separation of signal types, which neglects the possibility that thresholds may be nonlinear, in single or multiple dimensions, or more recently (ii) a variety of clustering algorithms to identify groups of points that are similar to each other in multiple dimensions [9,10,13–17]. The use of clustering algorithms reduces subjectivity associated with cluster identification and makes analyses repeatable. We have adapted this approach to scale easily across large SFCM datasets using limited computing resources.

Estimating the biovolume of cells and colonies is an important goal in the phytoplankton, which are responsible for nearly half of global primary production [18]. Cell biovolume is also a major determinant of how individuals interact with their abiotic and biotic environment. Called a ‘master trait’, biovolume influences nutrient-uptake rates, nutrient quotas, predator avoidance, and low-light performance [19–21]. Additionally, distributions of biovolume can - in combination with cell density (typically estimated by flow cytometers) - provide us with accurate estimates of total phytoplankton community biovolume. This community biovolume is highly correlated with total phytoplankton *biomass*, the most important algal parameter characterizing water quality. Despite its importance, quantifying biovolume using FCM and SFCM remains a challenge because of the complex shapes that phytoplankton take [22]. Present approaches to estimating biovolume involve the use of a single scattering channel and are based on calibrated relationships with beads [9,23]; they therefore assume that phytoplankton cells possess similar scattering properties to these beads. However, scattering properties vary between phytoplankton taxa based on differences in cell wall, vacuoles, and internal cellular structure, necessitating the development of better methods for biovolume estimation.

Identifying the functional groups that phytoplankton belong to is necessary if we are to understand ecological patterns because members of these groups share similarities in biochemistry, edibility, toxicity and roles in biogeochemical cycles [21,24]. Although we follow convention by referring to ‘functional groups’, these groups are defined taxonomically (e.g. cyanobacteria, diatoms, chrysophytes, green algae, etc.), but at different levels in the taxonomic hierarchy (phylum Cyanobacteria as opposed to class Bacillariophyceae). The term ‘functional group’ retains its usefulness because the groups are paraphyletic: some members of the clades/taxonomic groups are no longer phytoplanktonic (e.g. plants exist within the clade of the green algae). Identification of functional groups by SFCM is possible because groups differ in their autofluorescent photosynthetic pigments [24], leading to differences in cell absorption and emission profiles [25].

The protocol that we present here (outlined in Fig. 1) enables the quantification of the cell densities and biovolumes of phytoplankton communities and their major constituent functional groups. It relies on three tools: (i) multiple training datasets that we generated, (ii) an unsupervised clustering algorithm (flowPeaks [26]), and (iii) a machine learning algorithm (random forests [27]). These tools are used for three distinct roles: (1) identifying important variables for clustering, (2) classifying points into clusters, and (3) estimating cell biovolumes.

**Fig. 1.**
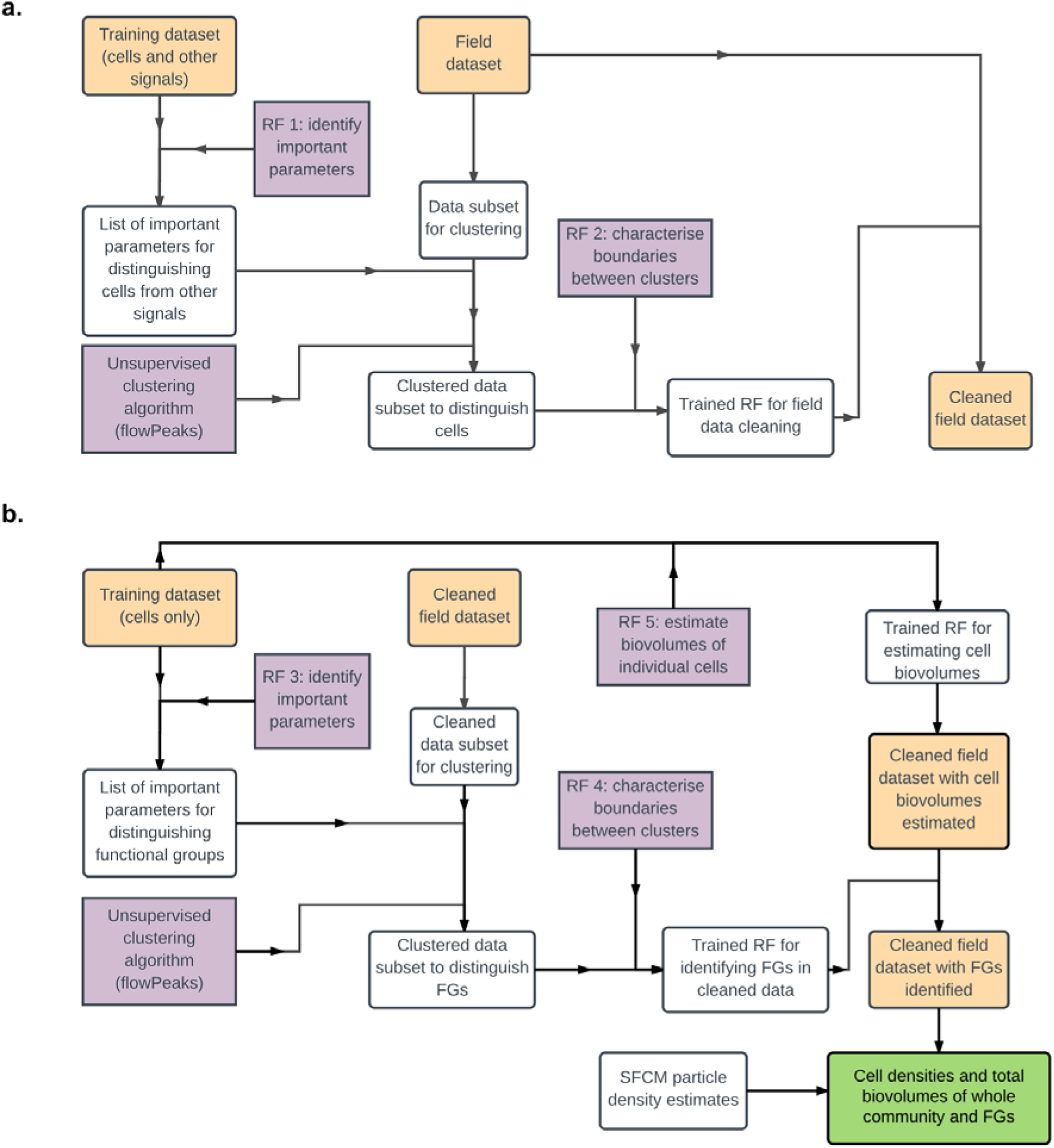
Flow chart illustrating the method we describe. We have divided this into two parts for clarity: a) Data cleaning, and b) Cell density and biovolume estimation. We use two abbreviations: RF for ‘random forest’ and FG for ‘functional group’. Yellow boxes indicate datasets, purple boxes indicate algorithms, the green box indicates the final product obtained, and white boxes contain other operations, objects or data manipulations.

Our advance consists in coupling these together in a manner that enables the processing and analysis of large FCM and SFCM datasets without the use of specialised computing resources such as high performance computing facilities. Unsupervised clustering algorithms identify groups of points that are similar to each other in multiple dimensions [28,29]. Here we use one such algorithm (flowPeaks [26]) that has previously been shown to provide reasonable results when applied to phytoplankton populations [17]. Random forests are a machine learning tool composed of ensembles of decision trees that can be used to identify features that separate groups in high-dimensional data. Each component decision tree uses a random subset of 66% of the dataset, and a randomly selected subset of variables that are assessed for correlation strength at each node. By aggregating predictions from all decision trees, random forests limit the overfitting problems associated with the use of individual decision trees [27]. No assumptions of linearity are required, with the algorithm capable of identifying curved boundaries in multiple dimensions. Additionally, they can be used to rank the importance of variables by assessing the differences in error rate (or alternatively, tree node purity [27]) when individual variables are permuted across all component decision trees. While a number of machine learning approaches could be applied to FCM and SFCM data, random forests have the advantage of being flexible, reasonably fast, and robust, negating the need for specialized computing resources.

We rely on training datasets based on lab cultures to (1) identify the traits that are best at distinguishing live cells from other signals, (2) distinguish functional groups from each other, and (3) quantify how all measured SFCM traits relate to cell biovolume measured by microscopy. Briefly, we used the most important variables identified by a random forest (applied to the lab culture training dataset) in combination with the flowPeaks algorithm [26] to clean SFCM data and to identify functional groups of cells in the cleaned data. The data we used to test this protocol were collected from a natural lake community across several months of automated monitoring. We used two additional random forests trained on clustered data to first classify all points from 191 files containing >20 million data points into phytoplankton cells and other particles, and subsequently classify the cells into clusters corresponding to functional groups. We used a fourth random forest trained on our lab culture training dataset to estimate the biovolume of every phytoplankton cell based on all the measured SFCM parameters.

We describe this procedure in detail below and validate it using 191 microscopy measurements corresponding to the same depths and times of the SFCM measurements (Fig. 1). We include the complete SFCM dataset that we used to evaluate this method on the data repository Zenodo, accessible at https://doi.org/10.5281/zenodo.977772 [30]. We also include code for the entire analysis on Github, accessible at https://doi.org/10.5281/zenodo.999747 [31], implemented in the R software environment [32]. Associated microscopy data used in this paper may be found in the supporting information.

## Methods

### Overview of approach

Our approach is shown in flow chart form in Fig. 1. It involves 10 steps:

(1) Generation of a training dataset using lab cultures with known species identity, functional group identity, and mean cell biovolume (Table 1).

**Table 1.** Cultures used in biovolume estimation. Mean biovolume values of these cultures were estimated by microscopy as part of this study.

| Species | Functional group | Coloniality | Mean biovolume ($\mu\text{m}^3$) |
| --- | --- | --- | --- |
| *Synechococcus* sp. | Cyanobacterium | Unicellular | 13.4 |
| *Microcystis aeruginosa* | Cyanobacterium | Unicellular (in culture) | 17.2 |
| *Chlorella* sp. | Green alga | Unicellular | 28.7 |
| *Ankistrodesmus bibrainus* | Green alga | Unicellular | 31.6 |
| *Chroomonas* sp. | Cryptophyte | Unicellular | 60.8 |
| *Synura* sp. | Chrysophyte | Unicellular | 152.7 |
| *Cyclotella cf. glomerata* | Diatom | Unicellular | 226.2 |
| *Cryptomonas* sp. | Cryptophyte | Unicellular | 896.8 |
| *Asterionella formosa* | Diatom | 6 cells per colony, with rare exceptions | 1964.8 |
| *Anabaena flos-aquae* | Cyanobacterium | Long filaments with highly variable numbers of cells. | 15478.7 |

(2) Identifying the parameters that most accurately distinguish between live cells and all other signals jointly (a combination of bacteria, detritus, and electronic noise), using the training dataset.

(3) Identifying clusters of similar points in a subset of the field dataset based on the parameters identified in step 2.

(4) Classifying points from the complete field dataset into these clusters, and removing all points except for those from live cells (data cleaning).

(5) Identifying the parameters that most accurately distinguish between different functional groups, using the training dataset.

(6) Identifying clusters of similar points (corresponding to different functional groups) in a subset of the cleaned field dataset based on the parameters identified in step 5.

(7) Classifying all cells from the cleaned field dataset into these clusters.

(8) Training a machine learning algorithm (a random forest) to estimate the biovolumes of individual cells based on all measured SFCM parameters, using the training dataset.

(9) Using the trained random forest from step 8 to estimate the biovolumes of every cell in the cleaned field dataset based on all measured SFCM parameters.

(10) Estimating cell density and biovolume of the total phytoplankton community and of the major clusters (corresponding to functional groups), using the assigned clusters for each individual cell (from step 7) as well as their biovolume estimates (from step 9) and cell density estimates.

### Procedures

#### 1. Sampling campaign

Between August 17^th^ and October 26^th^ 2014, we collected SFCM measurements at 6 depths (1.0, 2.5, 4.0, 5.5, 7.0 and 8.5 m) every 4 hours in Greifensee, a eutrophic lake in Switzerland (47.35°N, 8.68°E), using the automated monitoring station Aquaprobe [4]. We also collected weekly microscopy samples at every depth within 1 hour of an SFCM measurement, obtaining a set of paired SFCM and microscopy measurements. A small number of SFCM samples were lost, leaving us with 191 paired samples [30].

Over this time period, we saw environmental changes of approximately 15°C in water temperature, and variation of an order of magnitude in dissolved phosphorus concentration and N:P ratio.

The Office of Waste, Water, Energy and Air (AWEL) of Canton Zürich provided permission for *in situ* monitoring.

#### 2. Microscopy protocol

##### 2.1. Microscopy sample collection and counting

Microscopy samples were collected manually using a Niskin bottle. The Niskin bottle collects a vertical column of water 46 cms in length, which we centred at each of the 6 depths sampled by SFCM (therefore, the microscopy samples average over a water sample 46 cms in length as opposed to 1 cm in length as in the case of SFCM). 300 mL of water was fixed using Lugol’s solution and stored in a brown glass bottle in the dark till measurement.

Cells were counted using a Utermöhl counting chamber (3 mL) under 200X and 400X magnification [33]. 40 fields were counted and identified to a species level in most cases, and to the genus level in the remaining cases. Every sample was measured under both magnifications, with different species counted in each case based on their size.

##### 2.2. Microscopy biovolume estimation

The total biovolumes of the whole community and of the individual functional groups were calculated by multiplying the cell densities of each species (quantified by microscopy as described in section 2.1) by their mean per-cell biovolumes, and summing appropriately. The mean per-cell biovolume values of each species were taken from an existing, unpublished database of measurements made on individual cells from the same lake, Greifensee (Buergi unpublished), using standard protocols. This biovolume database is distinct from the measurements presented in Table 1 (which only refers to the training dataset). We used this second biovolume dataset for estimating biovolumes of field communities because i) we do not possess cultures of most species in the lake, and ii) though cultures of most species may be obtained from culture collections, these would be less representative, being strains from different environments and adapted to laboratory conditions. In contrast, our dataset was made using measurements of the same species from the same lake in previous years. Therefore, any influence of local environmental conditions on biovolumes was accounted for in these biovolume measurements. Intraspecific variation and changes through time were not considered, but we believe this is likely to be a small source of error because intraspecific variation in cell biovolume is considerably lower than interspecific variation (which can be >7 orders of magnitude, [34]).

#### 3. SFCM protocol

##### 3.1. Instrument description

SFCM measurement was performed using the CytoSense (http://www.cytobuoy.com), which is designed to characterize the scattering and fluorescence of individual particles; here we use it to study phytoplankton cells. The instrument measures particles across a large proportion of the phytoplankton length range, between approximately 2 µm and 1 mm in length. Although this instrument is capable of taking images, it is unable to resolve small cells clearly (approximately <10 µm). As these small cells are abundant in natural systems such as freshwater lakes, it necessitated the development of the method we present here.

Particles cross two coherent 15mW solid-state lasers (488 nm and 642 nm). Cells absorb and scatter these wavelengths, and also fluoresce at longer wavelengths determined by their specific pigment composition. Scattering (forward and sideward; the latter is typically referred to as ‘side-scatter’ in the flow cytometry literature) is measured, as well as fluorescence at each of three channels hereafter referred to as Red (668 – 734nm), Orange (601 – 667nm) and Yellow (536 – 601nm). These channels are listed below, followed by a more detailed description:

Red fluorescence Channel Laser 1: 488: 701/33

Red fluorescence Channel Laser 2: 642: 701/33

Orange fluorescence Channel: 488: 634/33

Yellow fluorescence Channel: 488: 569/33

Fluorescence in the Yellow and Orange channels is exclusively stimulated by the 488nm laser, while fluorescence in the Red channel is stimulated by both lasers. Therefore, the Red channel is electronically deconvoluted into Red 1 (from the 488nm laser) and Red 2 (a weaker signal from the 642nm laser). These wavelength bands are not highly specific in terms of the pigments they target, but generally target chlorophyll-a (Red 1), phycocyanin (Red 1 and Red 2), phycoerythrin (Orange), and carotenoids and decaying pigments (Yellow).

Hereafter, names of parameters representing the different fluorescence channels are preceded by ‘*FL*’, the Red 1 channel is referred to as ‘*FL.Red*’ and the Red 2 channel as ‘*X2.FL.Red*’. Names of parameters representing the scattering channels are preceded by *‘FWS*’ (forward scatter; typically abbreviated FSC in the flow cytometry literature) and ‘*SWS*’ (sideward scatter or side-scatter’ typically abbreviated SSC in the flow cytometry literature).

The signal produced by each particle is a time series of measurements for each channel, which describe the variation in scattering and fluorescence over the length of the particle. This high-resolution time series describes a pulse for each channel (4 fluorescence and 2 scattering). These pulses may be highly irregular in shape (Fig. S1) and are therefore characterized using a number of parameters (see Table S1 for parameter descriptions).

##### 3.2. Instrument configuration

Internal flow rates were set at 2 μL.s^−1^. A trigger threshold of 99.7 mV on the sideward scatter channel was enforced; particles whose scattering did not rise above this level were therefore not recorded. Field samples measurements were terminated when 500 µL was measured or 9 minutes elapsed, whichever was earlier.

##### 3.3. Field sampling procedure

Every four hours, the sampling tube was automatically deployed to each of the 6 depths by the automated station (described in [4]). Water samples were pumped to a 250 mL sampling chamber at the surface through a tube with a 1-cm diameter opening, making these highly depth-specific measurements. The sampling chamber was flushed with water from the sampling depth four to five times over 2 minutes before the CytoSense collected a water sample of up to 500 µL for measurement.

##### 3.4. Generation of training datasets using lab cultures

Our training dataset for the random forest served 3 purposes: (1) to enable the identification of parameters that most strongly distinguish between live cells and other signals, (2) to enable the identification of parameters that most strongly distinguish between cells belonging to different functional groups, and (3) to train the algorithm to estimate cell biovolume based on all the measured CytoSense parameters.

We measured ten laboratory cultures belonging to multiple functional groups (Table 1). In clonal lab cultures, manual identification of live cells and other signals is straightforward (examples in Fig. S2). We generated a dataset containing approximately 1,200 measurements of live cells (manually identified as in Fig. S2) and 8,400 measurements of other signals, equally sampled from all species’ measurements. The proportion was chosen to approximately mirror the low proportion of live cells expected from field measurements.

We note that overall performance will likely improve if the training dataset is improved. This may be done by increasing the number of species, number of functional groups, range of sizes, and range of culture conditions.

##### 3.5. Data cleaning

###### 3.5.1. Identifying parameters that distinguish live phytoplankton cells from other signals

The CytoSense measures 94 parameters for every particle. However, due to the ‘curse of dimensionality’ problem for clustering algorithms, we chose to reduce the number of variables examined. We used a machine learning algorithm, a random forest [27], to identify the variables that most strongly distinguished between live cells and other signals. Most importantly for this purpose, random forests can be used to generate a ranking of variable importance based on the change in classification error rate when a particular variable is permuted across individual decision trees. For this and all subsequent uses of random forests, we used the R package *randomForest* [35].

Before analysis, we log-transformed all variables whose minimum value was > 0, because most variables exhibited highly skewed distributions that could prove challenging for subsequent analyses. Then, to avoid problems associated with multicollinearity, we removed one member of all pairs of variables in the laboratory dataset whose correlation coefficient was > 0.8 or < −0.8. We were left with 51 variables. We trained a random forest with 10,001 trees on this dataset and found that the 10 most important variables enabled a clear distinction between live cells and other signals (Fig. 2). In descending order of importance, they were: *FL.Red.Range*, *X2.FL.Red.Range*, *X2.FL.Red.Gradient, FL.Red.Number.of.cells, X2.FL.Red.Last, FWS.Range*, *FL.Red.Fill.factor, X2. FL.Red.Fill.factor, FL.Yellow.Range*, and *FL.Orange.Range*. We note that the specific number of variables used is a feature of the dataset being considered, and readers are advised to explore using differing numbers of variables to identify the optimal number for their data.

**Fig. 2.**
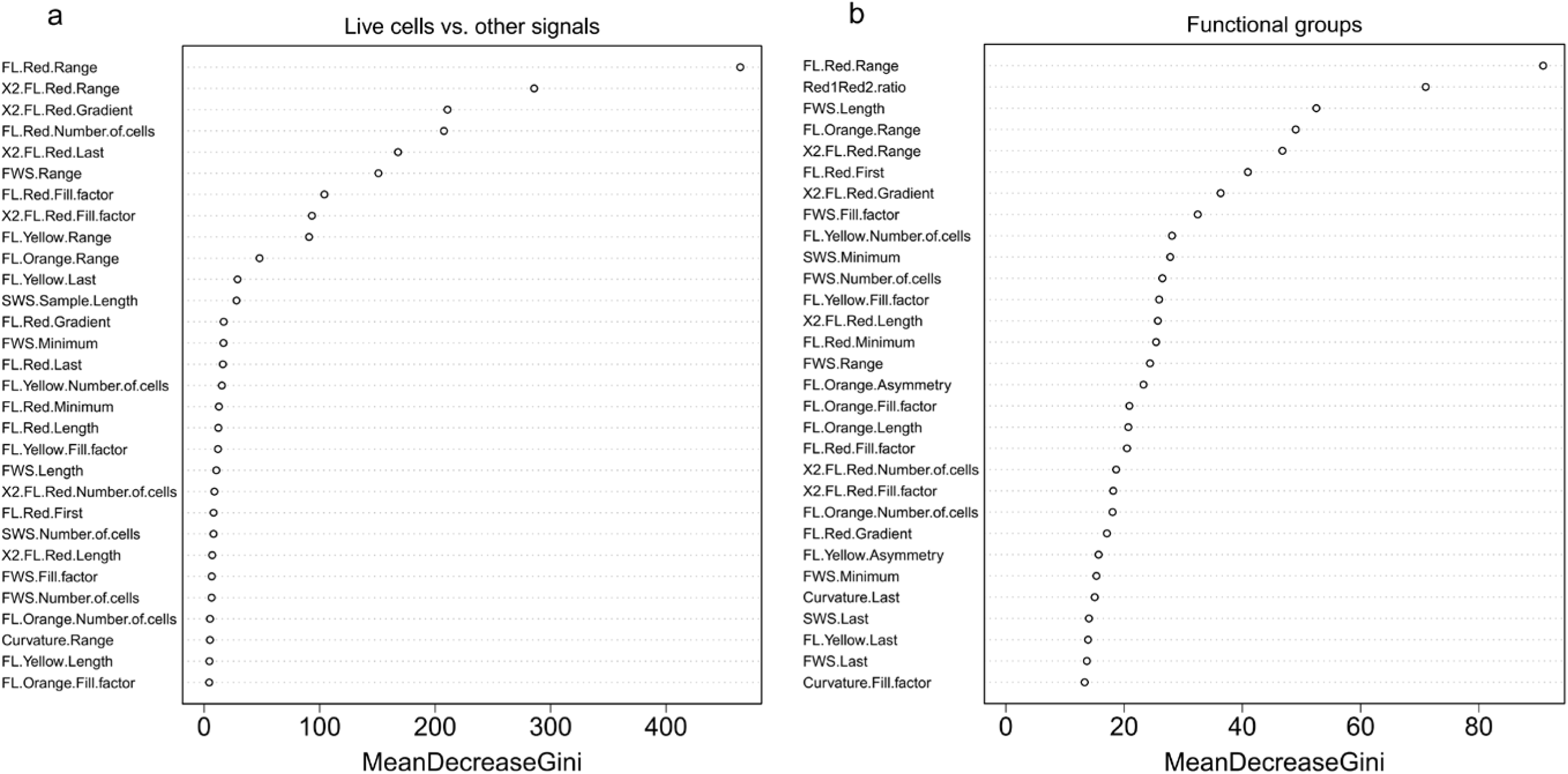
Importance of different traits in distinguishing between particle types, as determined by random forests applied to laboratory training data (Fig. S2). Variable importance is assessed using a Gini coefficient that measures the change in classification error when a variable is omitted [27]. The 30 most important variables are shown here. a) Variables that most strongly distinguish between live cells and other signals (electronic noise, detritus and bacterial cells). The top ten were used for unsupervised clustering of the raw field data. a) Variables that most strongly distinguish between major functional groups. The top eight were used for unsupervised clustering of the cleaned field data.

###### 3.5.2 Generating a random subset of the raw field data

The complete dataset of 191 measurements contained >20 million particles and could not be clustered simultaneously using standard computing resources. Furthermore, clustering each measurement separately would have led to the identification of different clusters as the community changed, limiting the possibility to make comparisons between measurements. We therefore generated a file containing an equal number of randomly selected data points from every measured sample, to distinguish between living phytoplankton cells and other signals. The data subset contained approximately 100,000 points which was both computationally tractable, and large enough to enable the identification of distinct clusters containing relatively low numbers of points that may be missed with smaller file sizes.

###### 3.5.3 Unsupervised clustering of the subset of raw data

We selected the 10 variables identified in section 3.5.1, log-transformed them and used the *flowPeaks* algorithm [26] to identify clusters of similar data points within the 100,000 point raw data subset. *flowPeaks* (implemented in the R statistical environment [32]) uses a k-means-based algorithm to first cluster the entire dataset into large numbers of small clusters, and then merges clusters based on density gradients. The smoothness of the density function is determined by tuneable parameters; we used values of 0.25 for *tol*, 0.05 for *h0*, and 2 for *h* [26] as these provided clusters that matched our expectations based on visual examination of 3D plots. However, the best parameter values may differ between datasets.

The algorithm identified 8 clusters, which we visually inspected using 3D plots (Fig. S3 and animated version in supplementary information, Table S2). From these plots, we identified 7 clusters that we expected to correspond to live phytoplankton cells; together they constituted 5% of the data. The remaining cluster corresponded to signals we were uninterested in, and cannot distinguish between: a combination of bacteria, dead phytoplankton, detritus, and electronic noise. Phytoplankton clusters were primarily characterized by high pigment fluorescence, particularly *FL.Red.Range* (a proxy for chlorophyll-a) (Table S2) and high forward scatter (FWS).

###### 3.5.4 Using random forests to classify raw data points and clean dataset

We used a second random forest [27,35] to classify all points in every sample into one of the eight clusters identified in the previous step using *flowPeaks* (code available at [31]). We trained the random forest with 1,001 trees on the clustered 100,000 point raw data subset, allowing it to identify nonlinear boundaries between the 8 clusters. The out-of-bag classification error rate of this random forest was <2%, indicating acceptably accurate performance in this classification step. When applying this method, this error rate (and the entire confusion matrix, or receiver operating characteristic curves) can be to evaluate whether the classifier performs acceptably well. We then used this random forest to classify all points in every sample into one of the 8 clusters identified earlier. Points identified as belonging to one of the 7 phytoplankton clusters were retained and the remainder discarded. The retained points from each sample were saved in separate files for further analysis.

#### 3.6 Functional group identification

We identified the functional groups that individual cells belonged to by essentially repeating the data cleaning procedure (section 3.5) on the cleaned data. There were minor differences in procedure that we note below.

##### 3.6.1. Identifying parameters that distinguish between phytoplankton functional groups

For this step, we used the lab culture training dataset containing 1,200 measurements of live phytoplankton cells only, belonging to five different functional groups – chrysophytes, cryptophytes, cyanobacteria, green algae, and diatoms (Table 1).

We then used a random forest to identify the variables that most strongly distinguished between these functional groups. As earlier, we log-transformed all variables whose minimum value was > 0 and removed one member of all pairs of variables in the laboratory dataset whose correlation coefficient was > 0.8 or < −0.8. We were left with 43 variables. We trained a random forest with 10,001 trees [35] and found that 8 variables enabled a clear distinction between cells belonging to different functional groups (Fig. 2). In descending order of importance, they were: *FL.Red.Range*, *Red1Red2.ratio*, *FWS.Length, FL.Orange.Range*, *X2.FL.Red.Range*, *FL.Red.First, X2.FL.Red.Gradient and FWS.Fill.factor*.

##### 3.6.2 Generating a random subset of the cleaned field data

To identify phytoplankton cells belonging to different functional groups, we generated a file containing an equal number of randomly selected data points from every cleaned data file, for a total of approximately 100,000 points.

##### 3.6.3 Unsupervised clustering of the subset of cleaned data

We selected the 8 variables identified in section 3.6.1, log-transformed them and applied *flowPeaks* [26] to identify clusters of similar data points. We used the same parameter values as earlier (*tol =* 0.25, *h0 =* 0.05, and *h* = 2). The algorithm identified 12 clusters, which we visually inspected using 3D plots (Figs. S4 and animated version in supplementary information, Table S3). We limited our subsequent analyses to the 4 clusters that formed >5% of the dataset (totalling >90% of the data), but note here that (i) functional groups may comprise multiple clusters and (ii) rarer functional groups are unassigned. The latter is not a limitation of the method; a more detailed laboratory comparison of the trait signatures of all functional groups would permit the assignment of every cluster to a functional group.

##### 3.6.4 Assignment of clusters to specific functional groups

Based on the fluorescence and scattering signals of these clusters and their relative abundance in the microscopy data, we assigned the 4 clusters to 4 functional groups. We note that our microscopy data showed that diatoms, dinoflagellates, euglenophytes, and desmids were almost entirely absent from the lake during the periods we sampled, leaving us with four well-represented functional groups – chrysophytes, cryptophytes, cyanobacteria, and green algae.

Cyanobacteria are characterised by a low *Red1Red2.ratio* value (indicative of high phycocyanin content), which was characteristic of Cluster 2 (Tables S3, S4). Our assignment of the remaining clusters is necessarily more speculative and would be improved by a broader sampling of the phenotypic diversity of these groups using lab cultures. Cluster 1, the most abundant group in the SFCM data, was designated as chrysophytes, the most abundant eukaryotic group in the microscopy counts. Cluster 3 was characterized by a low *X2.FL.Red.Gradient* and *FL.Red.First*, as well as an intermediate *Red1Red2.ratio*, indicative of cryptophytes. Cluster 4 was characterized by relatively high *Red1Red2.ratio* and *FL.Red.Range*, suggesting that they represent green algae. Note that in most cases other than *Red1Red2.ratio*, we did not have an *a priori* expectation relating to the importance of these variables in differentiating between functional groups. Therefore, we do not speculate on the reasons behind their importance here.

##### 3.6.5 Using random forests to classify cleaned data into trait clusters

We trained a random forest with 1,001 trees on the 100,000 point cleaned data subset, allowing it to identify nonlinear boundaries between the clusters identified in the previous stage. Out-of-bag error rates for this random forest were <2%, again suggesting acceptably low levels of classification error. We then used this random forest to classify all cells in the cleaned data files into one of the 12 trait clusters identified earlier. We recorded the cell densities of every trait cluster in every sample.

#### 3.7. Biovolume estimation

##### 3.7.1 Estimating the biovolumes of single cells and colonies

We trained a random forest with 10,001 trees on our lab culture training dataset (Table 1) to estimate the biovolumes of individual cells using all measured SFCM parameters. We used all the parameters and a training dataset with taxa from all the major functional groups in order to train a random forest that accounts for taxonomic differences in scattering and pigmentation properties. This approach was substantiated by our finding that *Red1Red2.ratio* (indicative of whether a cell was a cyanobacterium or a eukaryote) was the most important predictor of biovolume (Fig. S5) instead of scattering, which has previously been used in biovolume estimation of phytoplankton cells by SFCM [9,10]. Furthermore, the trained random forest had a model R^2^ (based on out-of-bag estimates) of 0.93 as opposed to a linear model based on the best single scattering variable (FWS.Length), which had an R^2^ of 0.50. We used this trained random forest to predict the biovolume of every individual cell in our cleaned field data based on all their SFCM parameters.

##### 3.7.2 Estimating the density of the whole community and major functional groups

We first quantified the cell density of each community by multiplying the particle concentration (estimated by the CytoSense) with the proportion of particles that were live cells in the sample. Subsequently, we estimated the cell density of each cluster (functional group) by multiplying the cell density of the whole community by the proportion of live cells in each sample belonging to the cluster.

##### 3.7.3 Estimating the total biovolume of the whole community and major functional groups

We estimated total biovolume of each community by multiplying the cell density estimated in section 3.7.2 with the mean biovolume of all cells in the community. Similarly, we estimated the biovolume of each cluster (functional group) by multiplying the estimated cell density of each cluster by the mean biovolume of all cells belonging to that cluster.

## Results and assessment

We assessed the accuracy of our data cleaning, functional group identification, and biovolume estimation procedures by using field data to compare SFCM estimates with microscopy estimates of (1) cell density at the whole-community level, (2) total biovolume at the whole-community level, (3) cell density of the major functional groups, and (4) total biovolume of the major functional groups.

### 1) Cell density of the phytoplankton community

SFCM estimates of whole-community cell density were strongly positively correlated with microscopy estimates (*r* = 0.73, *p* < 0.001, Fig. 3a). The slope of the regression between the two estimates is indistinguishable from the expected value of 1 (mean = 0.99, CIs: 0.86, 1.13) and significantly different from zero (p<0.001); the intercept is indistinguishable from the expected value of 0 (mean = −0.13, CIs: −0.65, 0.38). Quantitative estimates of cell density were extremely similar in both measurement methods. SFCM estimates were slightly lower on average, possibly due to (i) calibration of the SFCM inlet pump, leading to lower flow rates and consequently cell density estimates, (ii) difficulties in evaluating whether a cell was alive or dead in fixed microscopy samples, leading to overestimation of the microscopy cell density, or (iii) break-up of colonial organisms during preservation for microscopy. Whatever the underlying reason, the apparent difference is small, and variation in SFCM cell density estimates accurately captures variation in cell density by microscopy measurements.

**Fig. 3.**
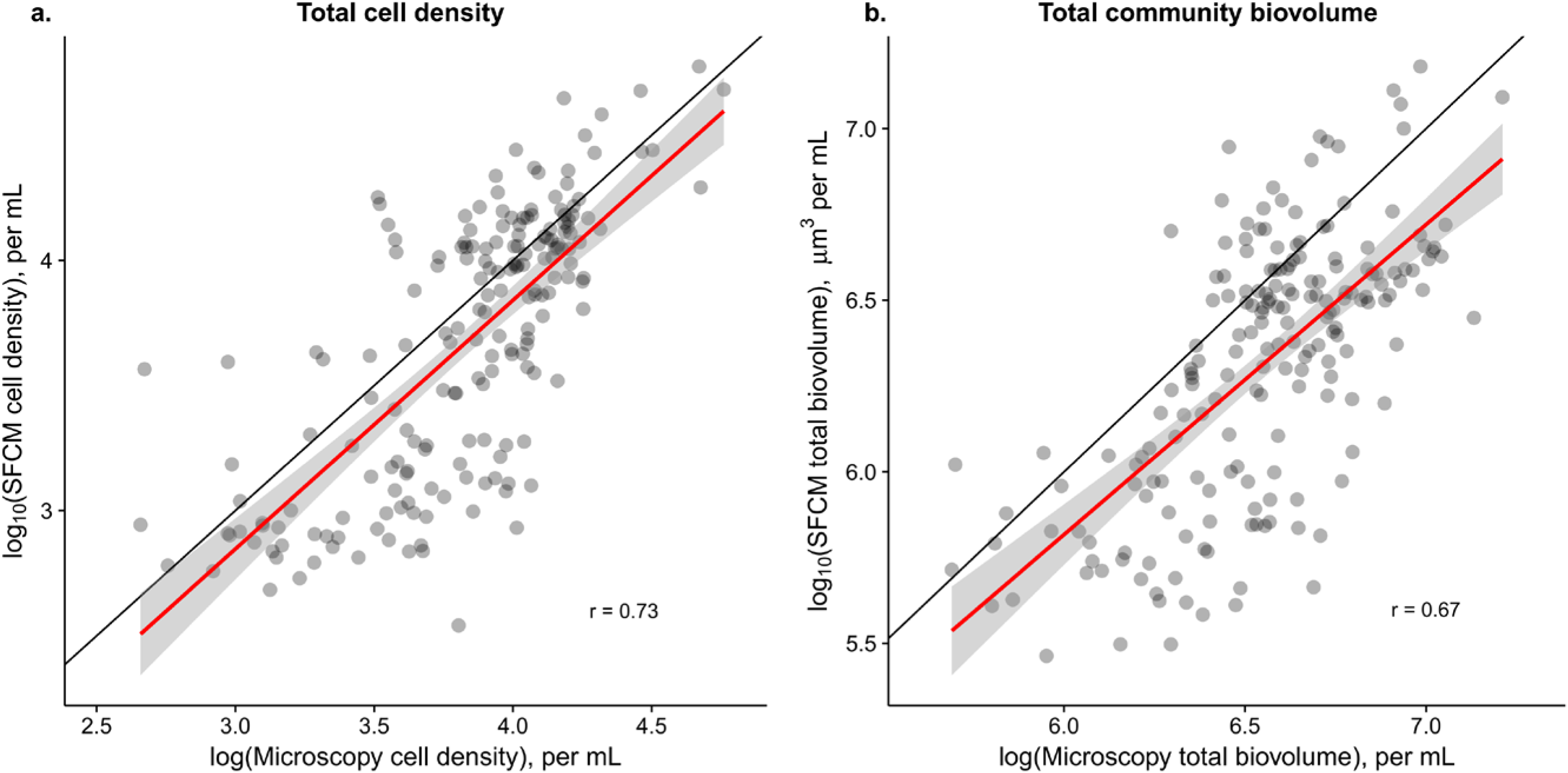
Estimates of whole-community cell density and biovolume by SFCM are strongly correlated with those by microscopy (on the log scale), and are quantitatively accurate. Each point represents a single depth-specific sample measured using both methods. The black line is the 1:1 line and the red line represents the regression relationship. a) Cell density estimates are strongly correlated (r = 0.73, p<0.001). b) Whole-community total biovolume estimates are strongly correlated (r = 0.67, p<0.001). The slope of the regression is indistinguishable from the expected value of 1 (mean = 0.90, CIs: 0.76, 1.05) and the intercept is indistinguishable from the expected value of 0 (mean = −0.40, CIs: −0.54, 1.33).

### 2) Total biovolume of the phytoplankton community

SFCM estimates of total community biovolume are strongly correlated with microscopy estimates (*r* = 0.67*, p <* 0.001, Fig. 3b). The slope of the regression is indistinguishable from the expected value of 1 (mean = 0.90, CIs: 0.76, 1.05) and significantly different from zero (p<0.001); the intercept is indistinguishable from the expected value of 0 (mean = −0.40, CIs: −0.54, 1.33). Quantitative estimates of total biovolume were similar in absolute magnitude over an approximately 30-fold range in biovolume, but appear to be slightly underestimated. This underestimation is partly driven by differences in cell density estimates (Fig. 3a), because density estimates are involved in total biovolume calculation. Differences in total biovolume are also influenced by our microscopy biovolume estimation procedure: estimates were generated by referring to a database of mean cell biovolumes and were not measured in the individual samples. Therefore, errors in our cell size database are propagated through to our estimates of total community biovolume by microscopy. However, intraspecific variation in cell biovolume is substantially lower than interspecific variation (which varies by >7 orders of magnitude [34]), and so we expect that this is at most a small source of error.

### 3) Cell density of the major functional groups

SFCM estimates of the cell density of major functional groups were moderately to strongly correlated with microscopy estimates (*r* ranges from 0.44 to 0.76*, p <* 0.001 in all cases, Fig. 4). Chrysophytes were the most abundant group overall and also show the strongest correlation and quantitative agreement between estimates. Cryptophytes and green algae show weaker correlations with slopes that were less than the expected value of 1, but they also exhibited relatively low variation over the sampling period, making it challenging to compare accurately. Cyanobacterial density estimates also showed a strong positive relationship, but this relationship was influenced by a small number of microscopy measurements in which no cyanobacterial cells were identified. These samples appeared to have been substantially overestimated by SFCM or underestimated by microscopy. This may partly be explained by the lower detection limit and consequently lower sampling error of SFCM measurements when compared to microscopy measurements, but may also be a result of cell degradation in microscopy samples which were not properly fixed. Though the quantitative estimates are substantially better if we exclude these points, the relationship remains reasonably strong even if we include them by assuming a cell density of 50% of the detection limit.

**Fig. 4.**
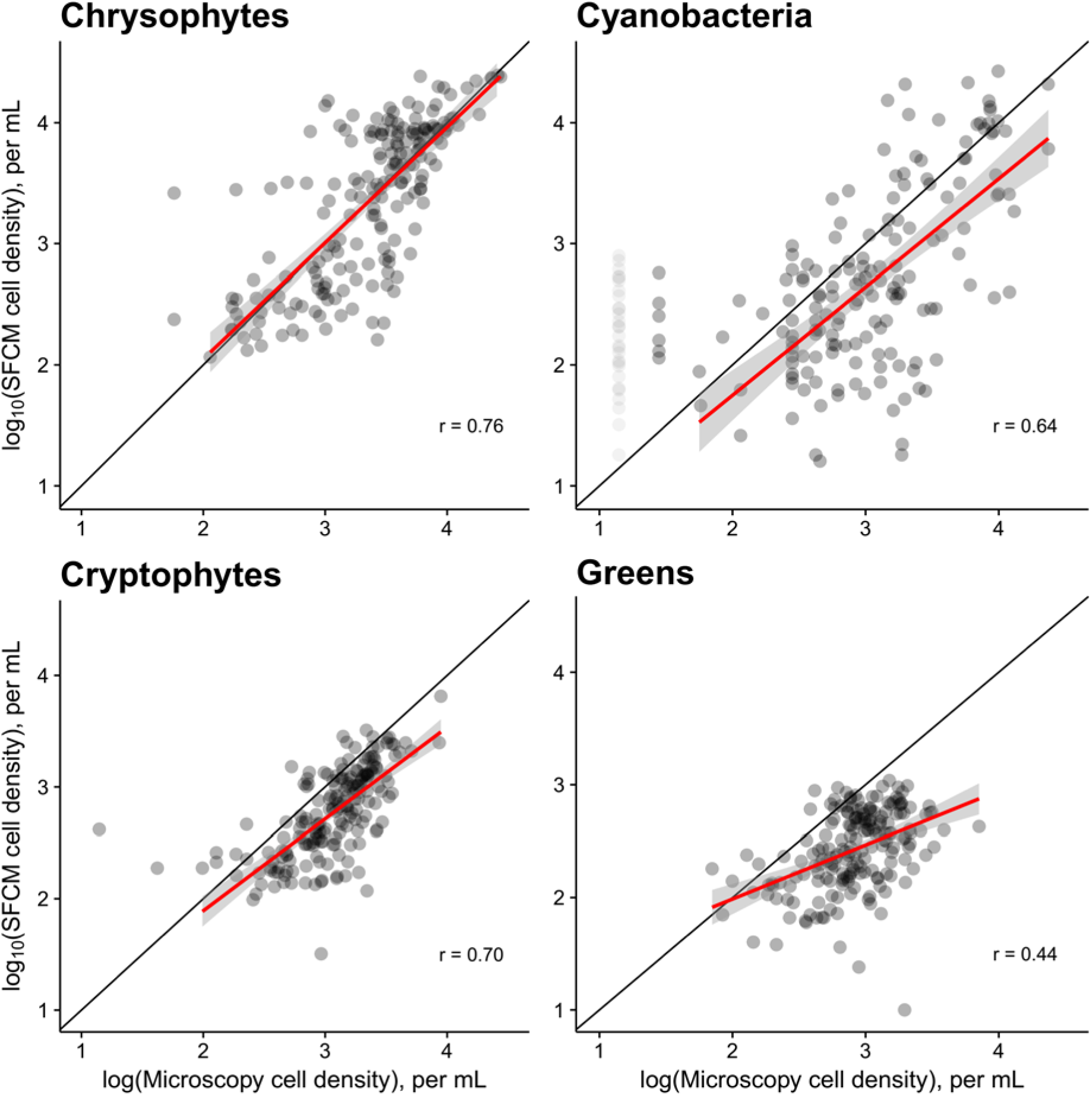
Estimates of cell density of four major functional groups by SFCM and microscopy are moderately to strongly correlated on the log scale (r between 0.44 and 0.76, p<0.001 in all cases). Each point represents a single depth-specific sample measured using both methods. The black line is the 1:1 line and the red line represents the regression relationship (with confidence bands). In a small number of microscopy measurements, no cyanobacteria were found. These points are plotted with high transparency assuming that the true concentration was half the detection limit of 28 cells per mL. A small number of high-leverage points were excluded from the regressions, but the effect of this exclusion on slopes and correlation coefficients is modest. These excluded points can be identified in the plots because the regression lines do not extend to them.

### 4) Total biovolume of the major functional groups

SFCM estimates of the total biovolume of major functional groups were moderately to strongly correlated with microscopy estimates (*r* ranges from 0.41 to 0.75*, p<*0.001 in all cases, Fig. 5). As with cell density, chrysophytes showed not just a strong correlation between estimates from the two methods, but also highly similar quantitative estimates. Cryptophytes and green algae exhibited a relatively small degree of variation and differed in absolute value, but were positively correlated in both cases. As in Fig. 4, cyanobacterial estimates were reasonably accurate, but a small number of measurements showed no cells in microscopy measurements even though the SFCM estimates were moderately high. Notably, slopes and correlation coefficients are relatively similar in both cell density and total biovolume. As cell density is used to estimate total biovolume, this suggests that the discrepancy is largely caused by the cell density estimates, and that mean cell biovolume of each group is being estimated accurately.

**Fig. 5.**
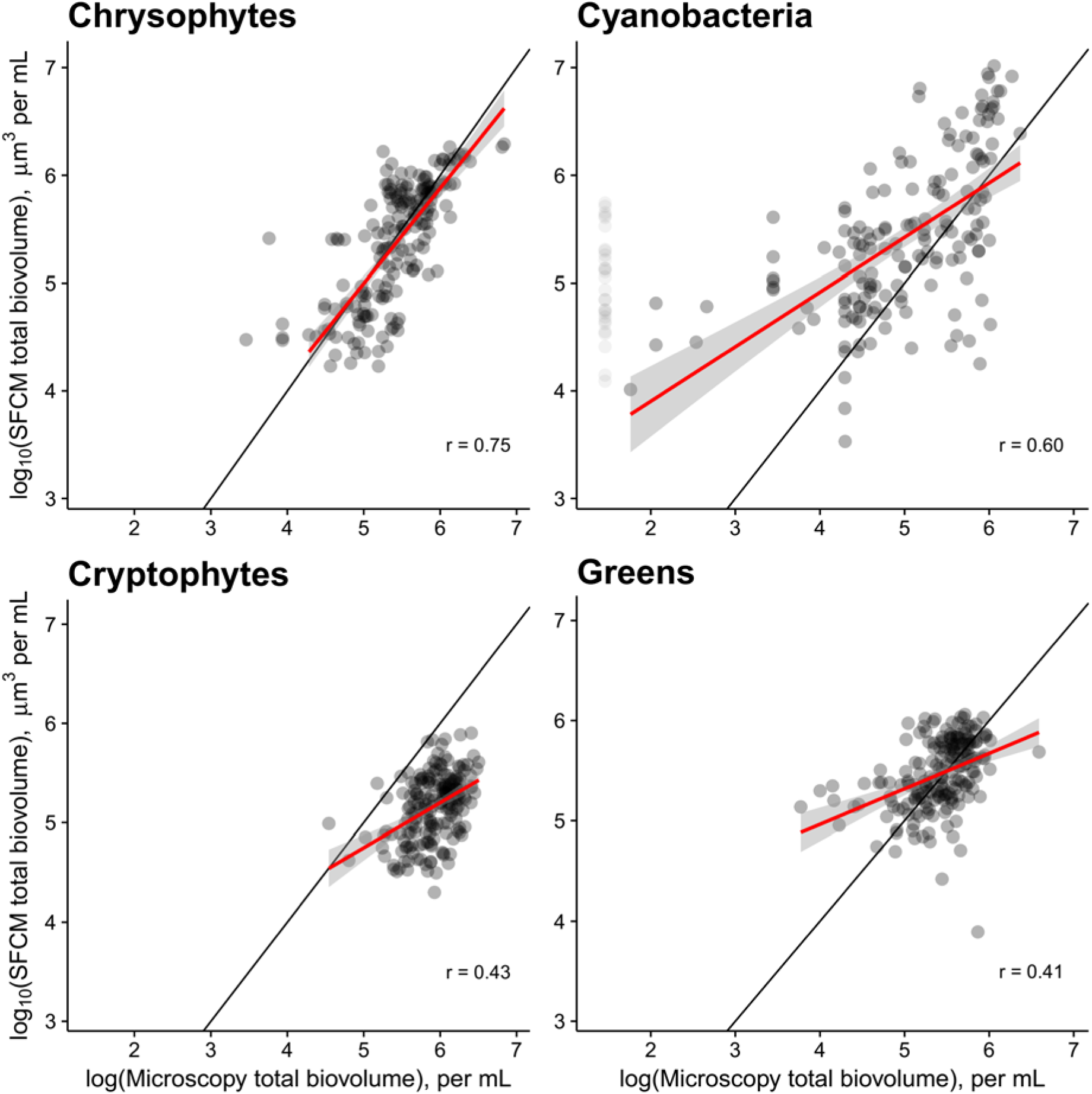
Estimates of total biovolume of four major functional groups by SFCM and microscopy are moderately well-correlated on the log scale (r between 0.41 and 0.75, p<0.001 in all cases). Each point represents a single depth-specific sample measured using both methods. The black line is the 1:1 line and the red line represents the regression relationship (with confidence bands). In a small number of microscopy measurements, no cyanobacteria were found. These points are plotted with high transparency assuming that the true concentration was half the minimum cyanobacterial biovolume detected. A small number of high-leverage points were excluded from the regressions, but the effect of this exclusion on slopes and correlation coefficients is modest. These excluded points can be identified in the plots because the regression lines do not extend to them.

We have demonstrated that this method produces estimates of cell density and biovolume that are reasonably well-correlated with estimates by microscopy, especially at the whole community level. These correlations may have been even stronger if microscopy samples could be collected at the same depth resolution as our SFCM measurements (we sampled a 46 cm-wide water column for microscopy and a 1 cm-wide water column for SFCM). And although we ignored less abundant clusters in this analysis because of limitations in the number of lab-measured taxa, our method can easily be expanded upon to include finer-scale variation and a wider range of functional groups.

Our protocol (Fig. 1) is composed of three parts that can each be improved upon with more data or improved algorithms, while following essentially the same protocol. The strong agreement between SFCM results and microscopy results from a complex natural community supports our claim. It improves on existing methods by (i) reducing the subjectivity associated with manual cluster designation, and (ii) increasing the scalability of FCM and SFCM analyses by enabling the study of datasets of arbitrarily large size using basic computing facilities. This scalability is achieved by coupling unsupervised clustering with machine learning techniques (which have frequently been used independently previously) and reducing the size of the major tasks through the use of training datasets and data subsets.

However, there are important limitations to our method:

1) The generation of training datasets is necessary and can be time-consuming.

Training datasets must contain representatives of all groups present in the system under study (with the possible exception of taxa that are rare both in terms of abundance and biovolume contribution). Functional groups are diverse and our training dataset for this study contains relatively few species from each cluster, potentially biasing biovolume estimates. However, this can be addressed by including more representative species from each group in the training dataset. Perhaps more importantly, this training exercise will need to be done independently for every instrument and possibly for changes in important instrument settings (such as photomultiplier tube gain) as well.

2) In addition to being time-consuming, the value of training datasets relies crucially on the assumption that traits measured are stable properties of individual cells, over time and space. If traits are stable, our approach should be applicable to conditions outside those under which training data are generated. The degree of stability in these traits is an empirical question that needs to be verified. Though we do expect variation to occur (e.g. nutrient starvation should lead to a decrease in pigmentation and therefore fluorescence), its magnitude is unknown. Based on our understanding of pigmentation and cell morphology (which influence fluorescence and scattering respectively), as well as our success here under variable conditions in a natural environment, we believe that traits are sufficiently stable to use this approach. However, if trait variation across environmental gradients is large, we would need to train machine learning algorithms with datasets that have been generated under conditions where there is a large degree of variability, or across multiple environments. This would improve performance by accounting for environment-dependent plasticity, which our random forest presently does not account for. Functional groups that are similar to each other in traits may also not be distinguished well (i.e. there may be misclassification) if this plasticity leads to changes in their trait distributions, or if they have contrasting trait-environment covariance patterns.

3) Many clustering algorithms have difficulties in identifying clusters that are highly nonlinear in multiple dimensions. Unsupervised clustering is an active area of research and future advances in clustering methods - especially density-based and neural network-based methods - may alleviate some of these problems [36]. However, every such algorithm has its own weaknesses and such improvements may come at the cost of other features (e.g. computational speed).

4) By clustering datasets with equal numbers of data points randomly selected from every time point, we might miss out on clusters that are present for very short periods of the time series. These clusters are poorly represented even if abundant for short time periods. This limitation is even more relevant if our approach is used to study datasets collected across space instead of time; clusters present at just one or two of a large number of sites may be difficult to detect. Addressing this issue will require careful sampling, but may also be aided by advances in clustering methods. A sequential clustering approach that re-clusters the data after the most abundant cluster is removed may also be useful in identifying these cases.

5) Our assignment of clusters to functional groups is at present provisional, except in the case of cyanobacteria, which are very strongly differentiated by their ratio of Red 1 to Red 2 fluorescence. This may be improved on by using instruments configured with more laser and detectors than the one used here, enabling more precise targeting of the pigments characterising different functional groups. In the absence of additional lasers and detectors, the problem may be mitigated by using larger training datasets with more representative species from each functional group.

Future work may therefore most profitably expand on our work by using a much broader training dataset to train machine learning algorithms. Capturing the entire range of functional groups and a broad sampling of the phenotypic variation within these groups would enable the classification of cells into functional groups (and potentially lower level taxonomic categories) without the need for unsupervised clustering.

## Discussion

Scanning flow cytometry has the potential to enable the high-frequency monitoring of natural phytoplankton communities in real time, improving our ability to understand their ecology [4,9]. For some ecological questions, such as those relating to the influence of short-term environmental changes on populations and communities, they are (along with imaging flow cytometers) the only feasible measurement tool at present. Tools to automate the analysis of FCM and SFCM data will greatly increase the utility of these instruments, increasing our ability to understand the drivers of high-frequency dynamics in natural communities.

A very large number of unsupervised clustering algorithms exist with differing strengths and weaknesses [28,29,37,38], and we did not attempt to compare the efficacy of different algorithms here. Instead, we use an algorithm (flowPeaks [26]) that has previously been applied to phytoplankton populations [17] and provides rapid, reasonably accurate results using relatively limited computational resources. More sophisticated algorithms that are capable of identifying points that do not below to any clear cluster (‘noise’) such as DBSCAN [39] may improve on this, but they generally require computationally expensive calculations, including the distance between all pairs of points to be clustered. Our approach is flexible and alternative algorithms can be substituted here if found to be more suitable.

A more general problem that our approach bypasses is that unsupervised clustering of high-dimensional SFCM data suffers from the ‘curse of dimensionality’. Output is less reliable as dimensionality increases, and visualisation of clusters also becomes difficult beyond three dimensions [40]. Therefore, there is a strong need to reduce the dimensionality of the problem for both reliable clustering and for visualisation. This may be done by two methods: (i) we could use a dimension-reduction algorithm such as principal components analysis, or (ii) we may limit the number of dimensions that we choose to cluster. We followed the second approach here because the former suffers from the weakness that the major principal components may not be meaningful in distinguishing between the signals we are interested in; in other words, there may be more variation in unimportant dimensions than in important ones. Although machine learning techniques have previously been applied to FCM data [12,41–43], our work differs in using it for tasks beyond classification. We have also used machine learning to predict biovolumes of individual cells, and to identify the phenotypic features that most reliably distinguish between particle types (Figs. 1, 2). When used in combination with unsupervised clustering, it constitutes a powerful and flexible tool in processing large FCM and SFCM datasets. In principle, a machine learning algorithm trained on lab cultures may be able to identify trait clusters in field data. However, in practice we need to couple this with unsupervised clustering because trait cluster boundaries differ between lab and field communities. This is because (i) natural communities usually contain species that are not in the training dataset, and (ii) changes in environmental conditions may lead to plastic changes in fluorescence and scattering, shifting the boundaries between clusters in multidimensional space.

Our method provides a repeatable, semi-automated protocol with which to process large FCM and SFCM datasets using readily available computing resources and free software tools (the R software environment and packages *randomForest* and *flowPeaks*). Our work integrates existing tools - lab measurements, machine learning, and unsupervised classification - from multiple fields to improve the speed, repeatability, and robustness of SFCM analysis.

We believe that this protocol will greatly increase the ease and speed with which large SFCM datasets can be processed and analysed. In the future, automated (or semi-automated) analytical tools such as these may enable the monitoring of water bodies for harmful algal bloom development in real time, through classification of cells based on previously developed training datasets. Cyanobacteria, which are responsible for freshwater harmful algal blooms, possess fluorescence traits that were especially easy to distinguish with our instrument configuration. Therefore, the identification and quantification of this taxon is simple. Examination of a broader range of cyanobacteria may allow us to identify features that distinguish between potentially toxic and non-toxic cyanobacterial taxa, improving the scope and accuracy of water quality monitoring efforts. However, this protocol will also improve our ability to study fundamental aspects of the physiology, ecology and evolution of phytoplankton communities, by enabling their monitoring on time-scales orders of magnitude lower than is presently feasible using microscopy methods.

## Acknowledgements

This work was supported by the Swiss National Science Foundation project CRSII2_147654, “In-situ Sensing Tools for Understanding Rapid Microscale Plankton Dynamics”. We would like to thank M. Stravs and L. Nizzetto for providing phytoplankton cultures, Esther Keller for microscopy analysis and Hannele Penson for assistance with field work, and Michael Kehoe for advice related to machine learning techniques. The Centre for Ocean Life is supported by the Villum Foundation.

## Statement of author contributions

MKT and FP conceived of the method with input from SF. FP, SF, MR & MKT collected the field data. MR maintained lab cultures, collected lab SFCM measurements and all microscopy measurements. MKT wrote the manuscript with substantial input from FP, SF and MR.

## References

1. Hammes F, Egli T. Cytometric methods for measuring bacteria in water: Advantages, pitfalls and applications. Anal Bioanal Chem. 2010;397: 1083–1095. doi:10.1007/s00216-010-3646-3

2. Sosik HM, Olson RJ, Armbrust EV. Flow cytometry in phytoplankton research. In: Suggest DJ, Prášil O, Borowitzka MA, editors. Chlorophyll a Fluorescence in Aquatic Sciences: Methods and Applications. Dordrecht: Springer Netherlands; 2010. pp. 171–185. doi:10.1007/978-90-481-9268-7

3. Wang Y, Hammes F, De Roy K, Verstraete W, Boon N. Past, present and future applications of flow cytometry in aquatic microbiology. Trends Biotechnol. 2010;28: 416–424. doi:10.1016/j.tibtech.2010.04.006

4. Pomati F, Jokela J, Simona M, Veronesi M, Ibelings BW. An automated platform for phytoplankton ecology and aquatic ecosystem monitoring. Environ Sci Technol. 2011;45: 9658–65. doi:10.1021/es201934n

5. Arnoldini M, Heck T, Blanco-Fernández A, Hammes F. Monitoring of dynamic microbiological processes using real-time flow cytometry. PLoS One. 2013;8: e80117. doi:10.1371/journal.pone.0080117

6. Hunter-Cevera KR, Neubert MG, Solow AR, Olson RJ, Shalapyonok A, Sosik HM. Diel size distributions reveal seasonal growth dynamics of a coastal phytoplankter. PNAS. 2014;111: 9852–9857. doi:10.1073/pnas.1321421111

7. Besmer MD, Weissbrodt DG, Kratochvil BE, Sigrist JA, Weyland MS, Hammes F. The feasibility of automated online flow cytometry for in-situ monitoring of microbial dynamics in aquatic ecosystems. Front Microbiol. 2014;5: 1–12. doi:10.3389/fmicb.2014.00265

8. Dubelaar GBJ, Gerritzen PL. A Step Forward towards Using Flow Cytometry in Operational Oceanography. Sci Mar. 2000;64: 255–265.

9. Pomati F, Kraft NJB, Posch T, Eugster B, Jokela J, Ibelings BW. Individual cell based traits obtained by scanning flow-cytometry show selection by biotic and abiotic environmental factors during a phytoplankton spring bloom. PLoS One. 2013;8: e71677. doi:10.1371/journal.pone.0071677

10. Pomati F, Nizzetto L. Assessing triclosan-induced ecological and trans-generational effects in natural phytoplankton communities: a trait-based field method. Ecotoxicology. 2013;22: 779–94. doi:10.1007/s10646-013-1068-7

11. Dubelaar GBJ, Jonker RR. Flow cytometry as a tool for the study of phytoplankton. Sci Mar. 2000;64: 135–156. doi:10.3989/scimar.2000.64n2135

12. Sosik HM, Olson RJ. Automated taxonomic classification of phytoplankton sampled with imaging in-flow cytometry. Limnol Oceanogr Methods. 2007;5: 204–216. doi:10.4319/lom.2007.5.204

13. Lo K, Brinkman RR, Gottardo R. Automated gating of flow cytometry data via robust model-based clustering. Cytom Part A. 2008;73: 321–332. doi:10.1002/cyto.a.20531

14. Aghaeepour N, Nikolic R, Hoos HH, Brinkman RR. Rapid cell population identification in flow cytometry data. Cytom Part A. 2011;79A: 6–13. doi:10.1002/cyto.a.21007

15. Malkassian A, Nerini D, Van Dijk MA, Thyssen M, Mante C, Gregori G. Functional analysis and classification of phytoplankton based on data from an automated flow cytometer. Cytom Part A. 2011;79 A: 263–275. doi:10.1002/cyto.a.21035

16. Glüge S, Pomati F, Albert C, Kauf P, Ott T. The challenge of clustering flow cytometry data from phytoplankton in lakes. In: Mladenov V, Ivanov PC, editors. Nonlinear Dynamics of Electronic Systems: 22nd International Conference, NDES 2014, Albena, Bulgaria, July 4–6, 2014 Proceedings. Springer International Publishing Switzerland; 2014. pp. 379–386.

17. Hyrkas J, Clayton S, Ribalet F, Halperin D, Virginia Armbrust E, Howe B. Scalable clustering algorithms for continuous environmental flow cytometry. Bioinformatics. 2015;32: 417–423. doi:10.1093/bioinformatics/btv594

18. Field CB, Behrenfeld MJ, Randerson JT, Falkowski PG. Primary production of the biosphere: Integrating terrestrial and oceanic components. Science. 1998;281: 237–240.

19. Litchman E, Klausmeier CA. Trait-based community ecology of phytoplankton. Annu Rev Ecol Evol Syst. 2008;39: 615–639. doi:10.1146/annurev.ecolsys.39.110707.173549

20. Edwards KF, Thomas MK, Klausmeier CA, Litchman E. Allometric scaling and taxonomic variation in nutrient utilization traits and maximum growth rate of phytoplankton. Limnol Oceanogr. 2012;57: 554–566. doi:10.4319/lo.2012.57.2.0554

21. Edwards KF, Thomas MK, Klausmeier CA, Litchman E. Light and growth in marine phytoplankton: allometric, taxonomic, and environmental variation. Limnol Oceanogr. 2015;60: 540–552. doi:10.1002/lno.10033

22. Sun J, Liu D. Geometric models for calculating cell biovolume and surface area for phytoplankton. J Plankton Res. 2003;25: 1331–1346. doi:10.1093/plankt/fbg096

23. Foladori P, Quaranta A, Ziglio G. Use of silica microspheres having refractive index similar to bacteria for conversion of flow cytometric forward light scatter into biovolume. Water Res. 2008;42: 3757–3766. doi:10.1016/j.watres.2008.06.026

24. Reynolds CS. The ecology of phytoplankton. Cambridge University Press; 2006.

25. Steinberg CEW, Schäfer H, Siedler M, Beisker W. Ataxonomic assessment of phytoplankton integrity by means of flow cytometry. In: Seiler JP, Kroftová O, Eybl V, editors. Toxicology - From Cells to Man. Springer Berlin Heidelberg; 1996. pp. 417–434.

26. Ge Y, Sealfon SC. Flowpeaks: A fast unsupervised clustering for flow cytometry data via K-means and density peak finding. Bioinformatics. 2012;28: 2052–2058. doi:10.1093/bioinformatics/bts300

27. Breiman L. Random Forests. Mach Learn. 1999;45: 1–35. doi:10.1023/A:1010933404324

28. Jain AK, Dubes RC. Algorithms for clustering data. Prentice-Hall, Inc.; 1988.

29. Jain AK, Murty MN, Flynn PJ. Data clustering: A review. ACM Comput Surv. 1999;31: 264–323. doi:10.1145/331499.331504

30. Thomas MK, Fontana S, Reyes M, Pomati F. Dataset: Quantifying cell densities and biovolumes of phytoplankton communities and functional groups using scanning flow cytometry, machine learning and unsupervised clustering. 2017. doi:10.5281/zenodo.977772. https://doi.org/10.5281/zenodo.977772

31. Thomas MK. R Code: Cleaning and clustering SFCM data. 2017. doi:10.5281/zenodo.999747. https://doi.org/10.5281/zenodo.999747

32. R Core Team. R: A language and environment for statistical computing. Vienna, Austria: R Foundation for Statistical Computing; 2017.

33. Utermöhl H von. Neue Wege in der quantitativen Erfassung des Planktons. (Mit besondere Beriicksichtigung des Ultraplanktons). Verhandlungen der Int Vereinigung für Theor und Angew Limnol. 1931;5: 567–596.

34. Litchman E, Klausmeier CA, Schofield OME, Falkowski PG. The role of functional traits and trade-offs in structuring phytoplankton communities: Scaling from cellular to ecosystem level. Ecol Lett. 2007;10: 1170–81. doi:10.1111/j.1461-0248.2007.01117.x

35. Liaw A, Wiener M. Classification and Regression by randomForest. R News. 2002;2: 18–22.

36. Stoop R, Kanders K, Lorimer T, Held J, Albert C. Big data naturally rescaled. Chaos, Solitons and Fractals. 2016;90: 81–90.

37. Parsons L, Parsons L, Haque E, Haque E, Liu H, Liu H. Subspace clustering for high dimensional data: A review. ACM SIGKDD Explor Newsl. 2004;6: 90–105. doi:10.1145/1007730.1007731

38. Aghaeepour N, Finak G, Dougall D, Khodabakhshi AH, Mah P, Obermoser G, et al. Critical assessment of automated flow cytometry data analysis techniques. Nat Methods. 2013;10: 228–238. doi:10.1038/nmeth.2365

39. Ester M, Kriegel HP, J S, Xu X. A density-based algorithm for discovering clusters in large spatial databases with noise. KDD. 1996;96.

40. Newell EW, Cheng Y. Mass cytometry: Blessed with the curse of dimensionality. Nat Immunol. Nature Publishing Group; 2016;17: 890–5. doi:10.1038/ni.3485

41. Balfoort HW, Snoek J, Smiths JRM, Breedveld LW, Hofstraat JW, Ringelberg J. Automatic identification of algae: Neural network analysis of flow cytometric data. J Plankton Res. 1992;14: 575–589. doi:10.1093/plankt/14.4.575

42. Boddy L, Morris CW, Wilkins MF, Al-Haddad L, Tarran G a, Jonker RR, et al. Identification of 72 phytoplankton species by radial basis function neural network analysis of flow cytometric data. Mar Ecol Prog Ser. 2000;195: 47–59. doi:10.3354/meps195047

43. Boddy L, Wilkins MF, Morris CW. Pattern recognition in flow cytometry. Cytometry. 2001;44: 195–209.

